# Spread of New Mutations Through Space

**DOI:** 10.1101/2022.01.07.475395

**Authors:** Kyle Shaw, Peter Beerli

## Abstract

The terms population size and population density are often used interchangeably, when in fact they are quite different. When viewed in a spatial landscape, density is defined as the number of individuals within a square unit of distance, while population size is simply the total count of a population. In discrete population genetics models, the effective population size is known to influence the interaction between selection and random drift with selection playing a larger role in large populations while random drift has more influence in smaller populations. Using a spatially explicit simulation software we investigate how population density affects the flow of new mutations through a geographical space. Using population density, selectional advantage, and dispersal distributions, a model is developed to predict the speed at which the new allele will travel, obtaining more accurate results than current diffusion approximations provide. We note that the rate at which a neutral mutation spreads begins to decay over time while the rate of spread of an advantageous allele remains constant. We also show that new advantageous mutations spread faster in dense populations.

## 1. Introduction

We seek to better understand the inherent difference between discrete and continuous population genetics models. One such difference arises with the distinction between population density and population size. Specifically, how do these two parameters affect the tradeoff between selective advantage and random drift. In discrete models, these two terms are often used equivalently, with the common knowledge that elevated population size decreases the effect of drift, while increasing the strength of selective advantage (Wright, 1931, Fisher, 1923, Gillespie, 2004, Gravel, 2016). However, after adding the spatial component required by continuous models, population density is no longer equivalent to effective population size; it is now defined as total population size divided by geographic area. While these two parameters may be highly correlated, it is still possible to have a spread-out sparse population with a high population count, or a dense population with a low population size. We investigate the interaction between the parameters population size, population density, and geographical area and their effect on drift and selection in a continuous model.

We also seek to better understand gene flow within a population. In discrete models, the term gene flow is a synonym for migration between discrete demes (Slatkin, 1985). However, in more uniformly continuous models, it is more natural to view gene flow as the movement of a gene across the landscape through a single population (Bradburd and Ralph, 2019). We model and predict how quickly a new gene is expected to flow through the population.

Despite the plethora of recent papers incorporating the spatial component into the traditional discrete models (Barton et al., 2002, Bradburd and Ralph, 2019, Selle et al., 2020, Schoville et al., 2012, Haller and Messer, 2019), the inclusion of a geographical component in population genetics models is not a new idea. From the very beginning of population genetics models, researchers such as Sewall Wright and Ronald Fisher have included spatial components in some of their models (Wright, 1943, Fisher, 1937).

The first in-depth, spatially oriented, population genetics models were presented by Sewall Wright 1943. He developed the Isolation by Distance (IBD) method to account for the fact that the farther apart two individuals were geographically, the more distinct their genetic code would be. Wright also introduced the concept of neighborhood size, *N_w_* = 4*πρσ*^2^ with *ρ* representing density, and *σ* representing dispersal distance. Realizing that some species can travel farther than others within a lifespan, neighborhood size scales with both population density and dispersal distance. This parameter can be loosely described as the number of neighbors an individual can interact with. Wright’s neighborhood size has been used extensively to correlate expected genetic diversity with geographical distance (Wright, 1946). Malécot and others have developed similar models and extended these methods (Malécot, 1948, Maruyama, 1972, Nagylaki, 1974). However, the spatial component of population genetics has proven to be very difficult and has a very rocky history. In 1975, Felsenstein discovered a major contradiction in the three major assumptions of these models. It is impossible to maintain a uniformly distributed population with a constant population size while placing children near their mothers according to a probability distribution such as a Gaussian (Felsenstein, 1975, 1976).

At this point, many researchers abandoned the investigation of purely continuous models and focused their work on a variation of the n-island model known as the stepping stone model (Wright, 1931, Kimura and Weiss, 1964, Malécot, 1975, Rousset, 2004). The expectation was that in the limit as *n* grows large, these models would be great approximations for continuous models (Robledo-Arnuncio and Rousset, 2010). The stepping stone model was originally developed to model populations that lived on islands or behaved in a similar fashion such as frogs in ponds (Dobzhansky and Wright, 1941). Until today, many researchers, including Rousset have used a variation of this model known as the lattice model (Rousset, 2008). Indeed, many of these methods have provided valuable insights into several genetic patterns. Rousset (1997) estimates the neighborhood size of a continuous population using the slope of the F-Statistic as the number of islands in his stepping stone method varies. Despite the abundance of the n-island model continuous approximations, we should be wary of the results obtained in the limit as *n* approaches infinity. Battey et al. (2020a) has shown that in some populations, some statistics diverge from the continuous model, including Tajima’s D and Wright’s *F_is_*

Recognizing that the spatial aspect of natural selection is crucial to understanding evolution and choosing to avoid the mathematical difficulties of spatial population genetics models, many methods look for simple correlations between genetic discontinuities and geographic features (Holderegger and Wagner, 2008). Thus, the field of landscape genetics was born. There is a plethora of methods and statistical analysis tools available for biologists and ecologists seeking to determine whether a geographic landscape feature such as a river or a mountain range is correlated with a genetic pattern such as a genetical cline or discontinuity. Mantel (1967) developed a statistical correlation test between genetic data and geographic location. However, Raufaste and Rousset (2001) explain that that this method incorrectly assumes there is no auto-correlation between samples, and that the extended partial Mantel test (Smouse et al., 1986) only removes linear correlation. Principal components analysis (PCA) is often used in conjunction with the site frequency spectrum and then overlayed onto a geographic map (Patterson et al., 2006, Hanotte et al., 2002, Menozzi et al., 1978). Others use variations of Monmonier (1973) and Womble (1951) to map genetic discontinuities onto a geographic map (McRae, 2006, François et al., 2010, Petkova et al., 2015). Yet until recently, very little work has been done that went beyond simple correlation and intrinsically tied a spatial dimension into a population genetics model.

Eventually Doebeli and Dieckmann (2003) developed a biologically relevant, spatially continuous population model. Rather than enforcing an arbitrary population size which must be maintained each generation, they instead let population size emerge as a result of the model. Population size now becomes a stochastic variable that, given proper parameters, will center around an average. Within real populations the carrying capacity of the environment and the interaction between individuals gives rise to the actual size of the population in each generation. The carrying capacity variable represents the expected number of individuals you might expect to live in a unit area. Whether this is limited by food or predators is unimportant in this context. Density dependent selection is introduced as a way of maintaining uniform distribution across the landscape and preventing the population from exploding. Leimar et al. (2008) completed an in-depth analysis on several spatial competition kernels including Gaussian and determined that this model does indeed lead to a uniform distribution, thus avoiding the contradictions Felsenstein discovered. Haller and Messer (2019) developed the code to accommodate this forwards in time model in their simulation software SLiM, and it has been used for several interesting discoveries (Battey et al., 2020a,b, Harris et al., 2018, Champer et al., 2020).

Barton and collaborators have also developed a density dependent spatial model (Barton et al., 2002, 2010a). They have incorporated large scale extinction and recolonization events, which increase the correlation between genetic diversity and geographic distance and potentially makes the model more realistic. The mathematics of this model have been analyzed rigorously (Etheridge, 2008, Etheridge and Véber, 2012, Berestycki et al., 2009, Kelleher et al., 2016). The probability of identity by descent was recalculated within this framework used for the spatial Λ-Fleming-Viot coalescent (Barton et al., 2010b, 2013). In essence, they incorporate geographic distance into the coalescent. The time to coalescence depends on both geographic distance as well as genetic distance. Kelleher et al. (2013, 2014) have implemented this model with the python module discsim. Since extinction and recolonization events are not pertinent to our study, we use Haller and Messer (2019)’s software, SLiM, to model a simpler situation.

Rather than mapping a genetic map onto a geographic map as is done in landscape genetics or incorporating a spatial scale on a well-defined discrete model such as the coalescent, we use the SLiM software to model a continuous population. We show how using a continuous landscape population model changes some basic assumptions about how alleles flow through a population. We introduce new alleles and observe how they spread through space and time; exploring the wave of advance Fisher first described in 1937. His original partial differential equation 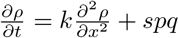 relates the allele frequency, *p*, of the new mutation at a location *x*, to a dispersal term, *k*, and selectional advantage, *s*, with *q* = 1 − *p*. See figure 1 for a visual of Fisher’s wave of advance. While this method was originally developed for new mutations, they have been successfully used to model the spread of several invasive species (Skellam, 1951, Reeves and Usher, 1989). However, in other cases they have failed to predict accelerating invasions which may arise due to genetic mutation increasing dispersal distance, or other geographic parameters (Urban et al., 2008). We use simulations to explore Fisher’s wave of advance of a new advantageous mutation under various population sizes and densities. We then develop a mathematical approximation for the speed at which a new allele, advantageous or neutral, will spread through a population.

**Figure 1:**
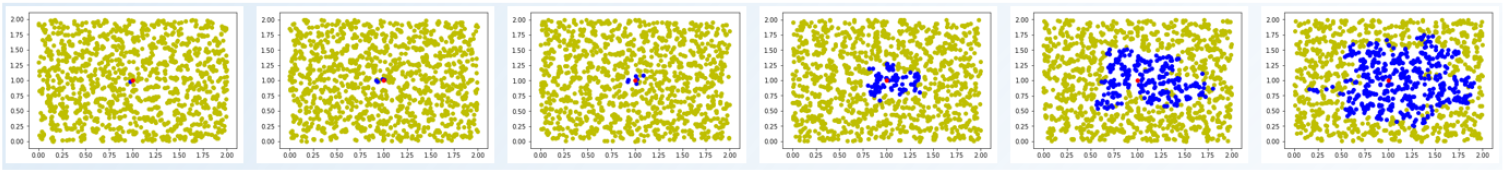
Snapshots of a typical simulation. Each diploid individual is plotted either as yellow or blue dot. Yellow represents a homozygote neutral whereas blue represents the presence if at least one advantageous allele in the individual. Moving to the right we jump several generations at each snapshot and see the wave of advance.

## 2. Material and methods

### 2.1. Simulations

A continuous population was modelled within a two-dimensional geographic space using the forward in time population genetics simulation software SLiM (Haller and Messer, 2019). Each generation consists of two main phases: first, reproduction and child placement, second, density dependent selection. In order to reduce computation time, each genome only consists of a single locus. At the beginning of the simulation all individuals are assigned two copies of the same neutral allele and an (*x, y*) coordinate. The individual closest to the center of the geographic space receives a mutation to one of their alleles with selectional advantage *s* ≥ 0 (*s* = 0 would be a neutral mutation).

Reproduction begins with mate selection. As a hermaphroditic diploid model, each individual has the opportunity to act as a mother and select a mate. Mates are selected from nearby geographic locations according to a 2D Gaussian distribution with parameters *u* = (*x_i_, y_i_*) and *σ* = (0.1, 0.1) where (*x_i_, y_i_*) is the location of the mother. The closer an individual is to the mother, the higher the likelihood of them being selected as the mate. If no potential mate can be found within 3*σ* then children will be produced via selfing. Once the parents are selected, the number of children is chosen according to a Poisson distribution with *λ* ≥ 1. Each child receives one allele from the mother and one from the father. The child is then placed into the population with a geographic location centered about the mother according to the same 2D Gaussian distribution (Figure 2). Once all the children are placed into the population, the parent generation is removed to simulate discrete generations.

**Figure 2:**
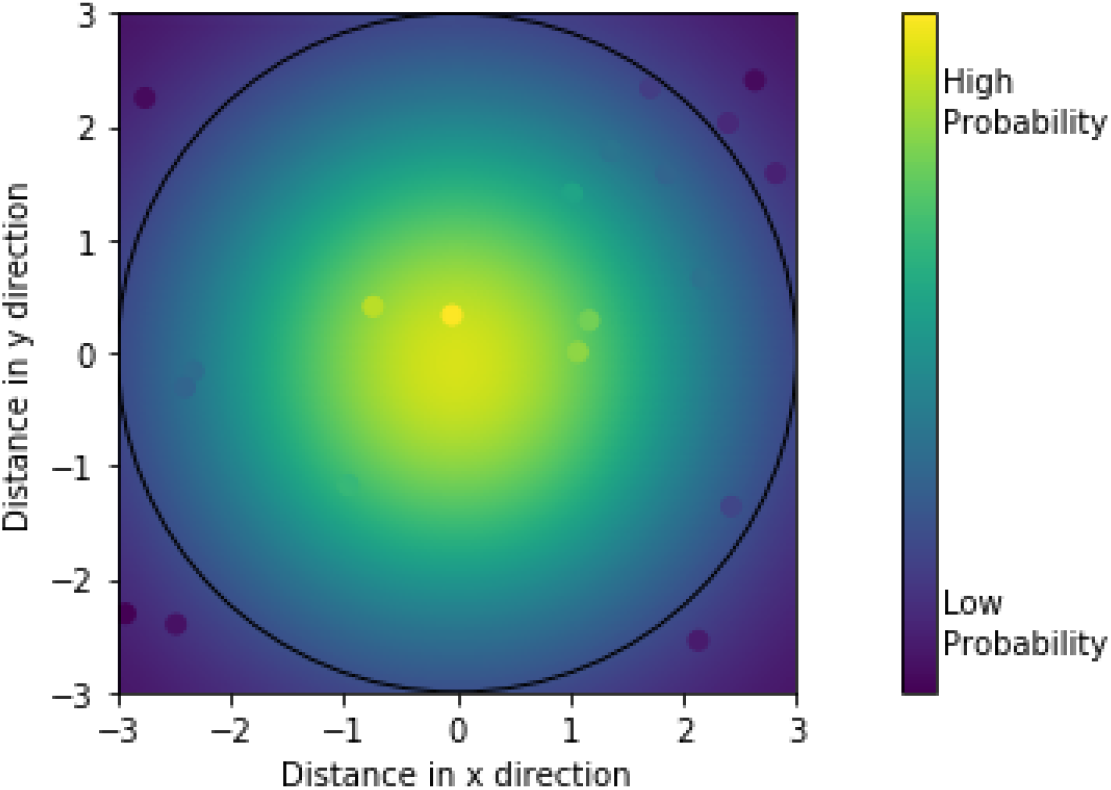
Example figure to demonstrate selective competition, mate choice probability, and probability of child location about the mother.

Density dependent selection is then incorporated into the probability of each individual surviving to adulthood. Since any environment can only sustain a finite population, we define the carrying capacity per unit area with the variable *K*. Carrying capacity is highly correlated with population density and can be used to approximate population size with the formula

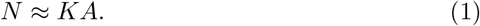

Where A is the total geographical area of the population. The probability of a child surviving to adulthood is calculated as the carrying capacity divided by the current local density of the children. Any value greater than 1 implies the carrying capacity is large enough to negate any competition felt by the current population. This probability is capped off at 1 since you can’t have greater than 100% probability of survival. To account for the fact that competition strengths vary with distance, the current local density of an individual is calculated as the summation of all competition strengths felt by that individual. We again use a 2D gaussian curve with parameters *u* = (*x_i_, y_i_*) and *σ* = (0.1, 0.1) to calculate these values. Letting *d_ij_* be the distance between individuals *i* and *j*, the probability of individual *i* surviving to adulthood is

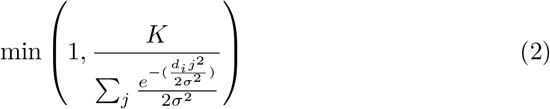

Replacing probability for competition strength in Figure 2 provides a visual of how competition strength varies according to distance (Doebeli and Dieckmann, 2003, Leimar et al., 2008, Haller and Messer, 2019).

Each generation, after selection, the distance is measured between the point of origin and each mutant. Over time these mutants will be found further and further from the point or origin. The maximum distance is viewed as the front edge of the wave of advancement as described by Fisher (1937). Since this process is stochastic and highly variable, over 10000 simulations were run for each set of parameters and averages calculated.

## 3. Theory/calculation

### 3.1. Mathematical Model

Random walks are often the starting point when dealing with models of dispersal. Consider the position of an individual and the positions of each ancestor/mother of that individual. Assuming each child is centered about the mother according to a Gaussian distribution, this sequence follows a random walk. We will call it the ancestral random walk. The dynamics of a large set of ancestral random walks are often modeled with diffusion PDE approximations (Skellam, 1951). These approximations can be quite accurate when dealing with a large number of independent random walks. We, however, are interested in a small number of somewhat correlated random walks. Using probability theory, we derive probability functions and expectations which are much more descriptive of reality.

We first model the introduction of a neutral allele. At any time *t* there will be *k* individuals with the new mutation. Each of these *k* individuals will have an ancestral walk back to the point of origin. We model the front edge of Fisher’s wave of advance as the expected maximum distance of those *k* walks. In the simplest case we have *k* = 1. The probability density function for the distance *r* after *t* generations can be written as

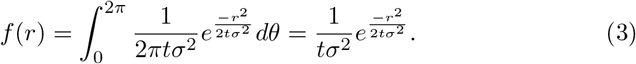

The expected distance this 2D Gaussian random walk travels in *t* generations

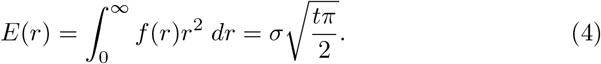

The expected speed the allele travels can be calculated as the derivative of equation 4, which evaluates to

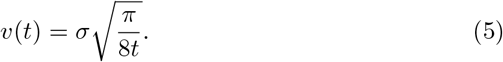

Letting *k* > 1 is more complicated as we need to calculate the expected distance the farthest individual will have travelled. The cumulative density function (CDF) of this maximum distance is most readily determined. The probability density function (PDF) can be calculated as the derivative of the CDF, then used to calculate the expected distance as before. Letting *r_i_* be the distance an individual *i* is found from the point of origin at time *t* with *i* = 1…*k*, we set *R* as *max*(*r_i_*). The CDF of *R* is

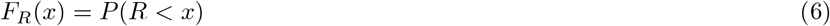

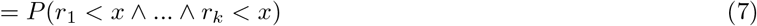

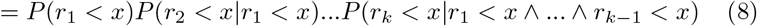

The probability that any *r_i_* < *x* given any other *r_j_* < *x* is a very complicated formula and requires either knowledge of the ancestral genealogy, or a summation over all possible genealogies which can be a rather large space. Thus, we make the rather dubious assumption of independence. When considering the effects this has on the equation, we notice that *P*(*r_i_* < *x*|*r_j_* < *x*) must always be greater *P*(*r_i_* < *x*), Thus this equation will be a lower bound. Since each walk is identically distributed this equation simplifies further

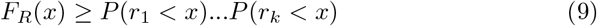

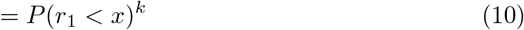

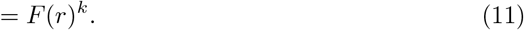

*F*(*r*) is the CDF for the distance of a single random walk which can be calculated as the integral of equation 3. The PDF of *R* can now be calculated as

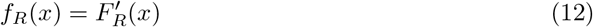

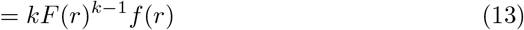

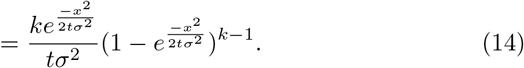

We now calculate the lower bound for the expected distance of the maximum distance of *k* random walks in *t* generations as

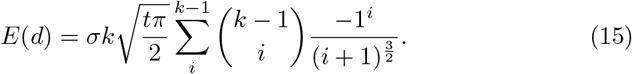

with the instantaneous rate of speed calculated as the derivative which gives

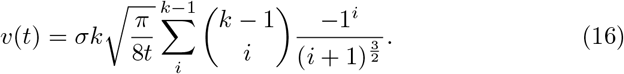

Equation 16 shows that the spread of neutral mutations will begin to slow over time. *k_t_*, or the number of descended individuals in generation *t* can be calculated according to a Poisson distribution conditional upon non-extinction.

Modelling the spread of an advantageous allele is more difficult. Given that the new mutation is beneficial, mutants on the leading edge of the wave will be competing with less fit individuals containing the ancestral allele. The mutants on the wave edge will have more children than any other individuals within the population. This creates a bias toward individuals with the new mutation travelling outward, thus increasing the correlation between random walks even further than that seen in the neutral case. If this bias scales with 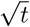 then the rate at which an advantageous allele spreads through a population remains constant rather than slowing down as it does in the neutral case.

Rather than modelling the spread of an allele with a random walk, we consider the probability of an individual possessing the advantageous allele given its distance from the point of origin. We then update this probability curve to match the expected values of the next generation after dispersal and then update it again to match the values expected after selection (Figure 3).

**Figure 3:**
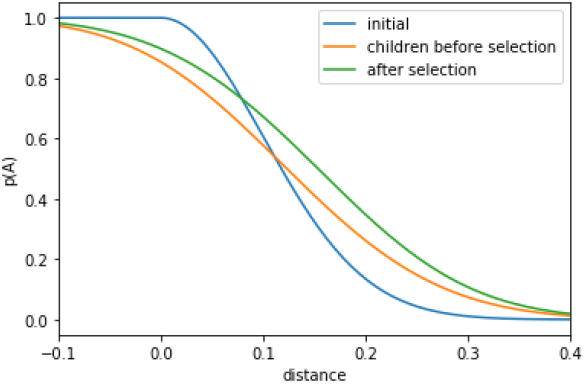
Probability of Advantageous allele across geographic distance. The x-axis is arbitrarily set with *x* = 0 as the trailing edge of the wave in the initial frequency graph.

Assuming that the speed of the wave of advance is constant, as will be shown later, we measure the distance this wave moves between each generation to calculate the speed of advance. Since we do not know what the probability curve looks like at any given *t* we use an initial guess for the probability curve, then recursively let the scattering of children, and the selection of individuals update the curve until is converges to the true probability curve. Once the wave shape converges, we can measure the distance the wave travels in each generation.

Let the initial probability of the advantageous allele be *I*(*x*). Then the probability curve after children dispersal *P_c_*(*x*) can be calculated as 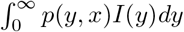. where *p*(*y, x*) is the probability that the parent of an individual at location *x* came from location *y* (Skellam, 1951). We assume children are centered about parents according to a Gaussian distribution so this would be 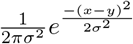. Thus, we obtain

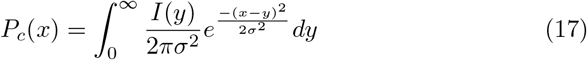

Selection will increase the probability of seeing the new advantageous mutation. There are several formulae for calculating the new probability frequency of an allele after one generation. The most popular formula can be found in Gillespie (2004)’s book and is

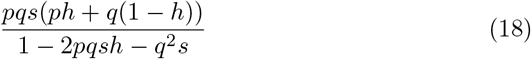

where *s* is the selectional advantage and *q* = 1 − *p* where *p* is the probability of being an advantageous allele, and *h* is the dominance ratio. This model assumes an infinite population size, which clearly doesn’t suite our needs. Of the other methods which calculate this new probability frequency given a finite population they either assume a fixed population size, which affects the probability calculations (Wright, 1946, Jensen and Pollak, 1969), or they make approximations using diffusion equations, often using the phraseology ‘‘Error is negligible if [population size] is large.” (Jensen, 1973, Kimura, 1964, Takahata et al., 1975).

Ewens (1963) finds that diffusion approximations are valid for populations that are reasonable. However, when dealing with only the population found within a small locale, the population sizes are clearly much smaller than Ewens would consider reasonable, especially when dealing with low density populations. Using probability theory, we calculate the exact probability density function of a new allele frequency given the previous allele frequency.

The expected frequency of an allele after selection is calculated as the summation of all possible frequencies weighted by their probability. In finite populations these can be calculated with a modified binomial distribution. Let the frequency of the advantageous allele before selection, at location *x* be *P_c_*(*x*) and let *N* be the expected population size with c being the number of children each individual has on average. Thus *Nc* is approximately the number of children before selection. Let *r_p_* be the probability of survival for the advantageous alleles, and *r_q_* be the probability of survival for the neutral alleles which are 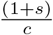 and 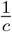 respectively. The Expectation of *P*(*x*) after selection is

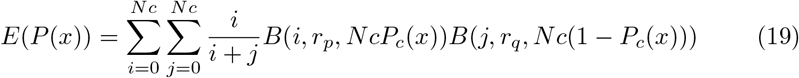

where *B*() is the Binomial probability function. We note that this formula is undefined when *i* and *j* are both 0, which would be the rare case where all individuals, both mutants and non-mutants, fail to survive the selection step. In the case of this local extinction event, the frequency of the mutant allele is uninteresting. Thus, we instead calculate the expectation of *p* conditional upon non-extinction of the local population. To calculate this, we skip the single event where both *i* and *j* are zero and divide our result by the remaining probability of

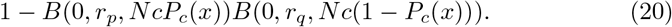

This formula has no closed form solution but can be calculated numerically. The expected change in *p* i.e., *p_new_* − *p_old_* for various population sizes is plotted in Figure 4.

**Figure 4:**
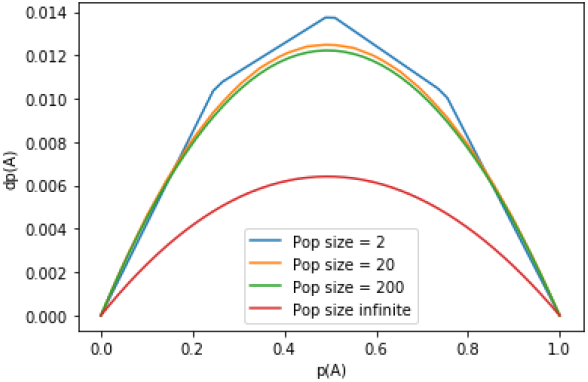
Expected change in allele frequency of an advantageous allele given each individual has two children, and a selectional advantage of 0.05

To summarize, we start with an initial probability curve *I*(*x*), then we recursively calculate *P_c_*(*x*) and *P*(*x*) after both dispersal and selection until the shape of *P*(*x*) and the distance it travels in each generation converges. These equations are given as

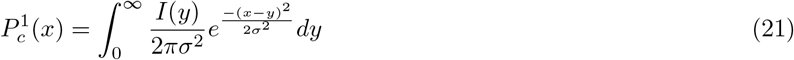

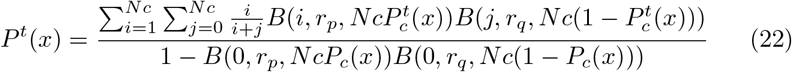

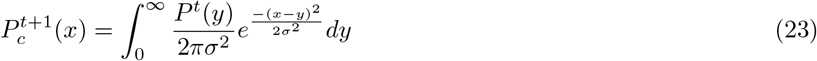

We calculated the probability curve and distance travelled of a population with Wright’s neighborhood size, *N_w_* = 2. In figure 5 we see that the distance travelled does converge despite the initial guess *I*(*y*) of the frequency of advantageous alleles as seen in figure 3.

**Figure 5:**
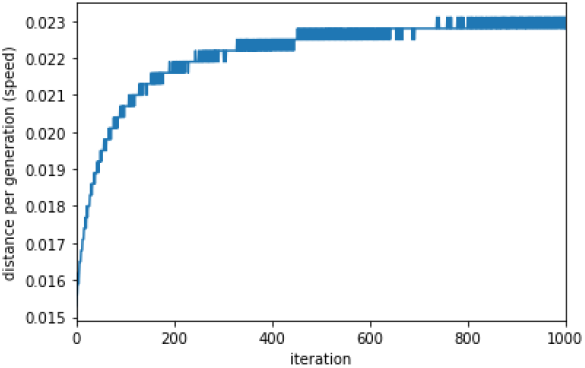
Distance that the wave travels in each generation. Converges to about 0.023 with *N_w_* = 2.

## 4. Results

Several basic results can be gleaned from animations of Fisher’s wave of advance. Since videos can be difficult to show in a static document, Figure 6 helps visualize what a wave of advance is at different times and in different conditions with several snapshots of various simulations. These simulation snapshots demonstrate some of the fundamental differences between dense and sparse populations that may not be as apparent when viewing the summary statistics or averages. An advantageous allele will saturate a dense population much more quickly than it will in a sparse population. The leading edge of the wave of advance is more circular and expands quicker than the more jagged edge found in the sparse population. The neutral mutation snapshots imply that population density plays a minor role in the spread of neutral mutations.

**Figure 6:**
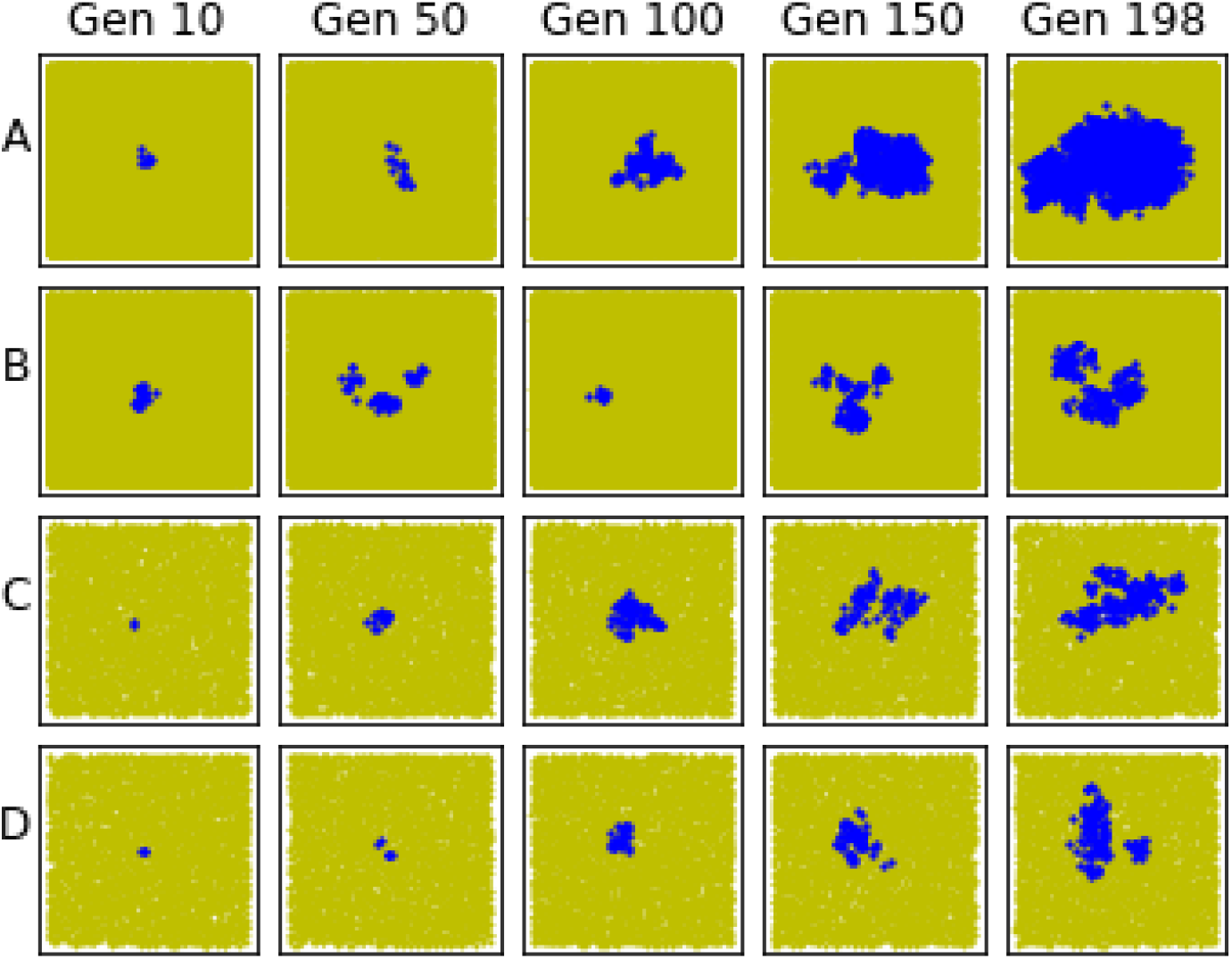
Snapshots of four typical simulations. Each column is a snapshot at the specified generation. All simulations were ended when a mutant allele approached the boundary (i.e. Generation 198). A: Dense population (K = 200) with an advantageous allele (s = 0.05). B: Dense population with a neutral allele (s = 0.0). C: Sparse population (K = 50) with an advantageous allele. D: Sparse population with a neutral allele.

After running 10,000 simulations for an advantageous allele and for a neutral allele, we plotted the distance travelled versus generation number. In Figure 7 we plot this distance travelled along with the calculated speed. The actual speed of the wave of advance is calculated using the derivative which is approximated with the centered difference formula.

**Figure 7:**
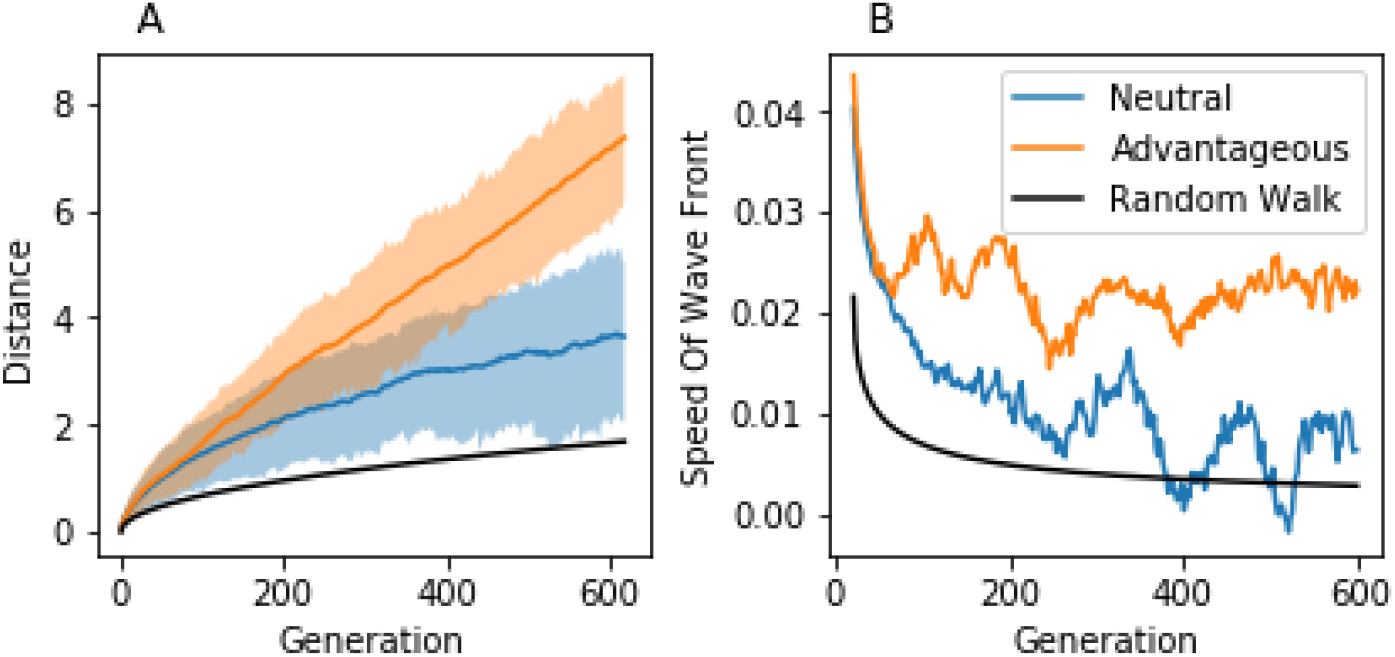
(A) Maximum distance that the new allele is found from the point of origin at each generation. For each simulation, 10,000 runs are performed, and the average is computed and displayed as a solid line. The shaded region about each line marks the range from the 10th to the 90th percentile; for the advantageous allele *s* = 0.05 and the neutral allele *s* = 0.0. The theoretical expected distance for a single random walk is also shown. (B) Speed of the plots on the left. Derivatives were calculated with the centered difference formula.

We note that initially both waves travel fairly quickly. In the first generation, children are guaranteed to move away from the point of origin. As time passes and the mutant’s descendants are found further from the point of origin, the probability of dispersing sideways or even back toward the point of origin increase. This slows the rate of spread over time; at least initially. Note that the speed of the wave of advance for the advantageous allele does not decline indefinitely but stabilizes at about 0.025. This constant speed causes the distance curve to appear linear with respect to time. Referencing equations 5 and 16, we predict that the distance graph of the neutral allele should scale with 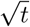, and the derivative should converge to 0 as *t* → ∞, which is indeed the case.

To see the effect of various densities, we ran 10,000 simulations for each density *K* = 25, 50, 100, 200, 300, 400, 500 on both an advantageous allele and a neutral allele (Figure 8). When considering an advantageous allele we notice that increasing population density will increase the speed of the travelling wave. The neutral allele, however; shows no dependence on population density. We find this quite remarkable. This implies that the spread of neutral mutations is affected only by the geographic distance a species covers and not by the population size or density. In fact, when calculating *k* for equation 16 we recognize that there is little dependence on the population size or density. The prediction from equation 16 is plotted alongside a simulated neutral allele in figure 9.

**Figure 8:**
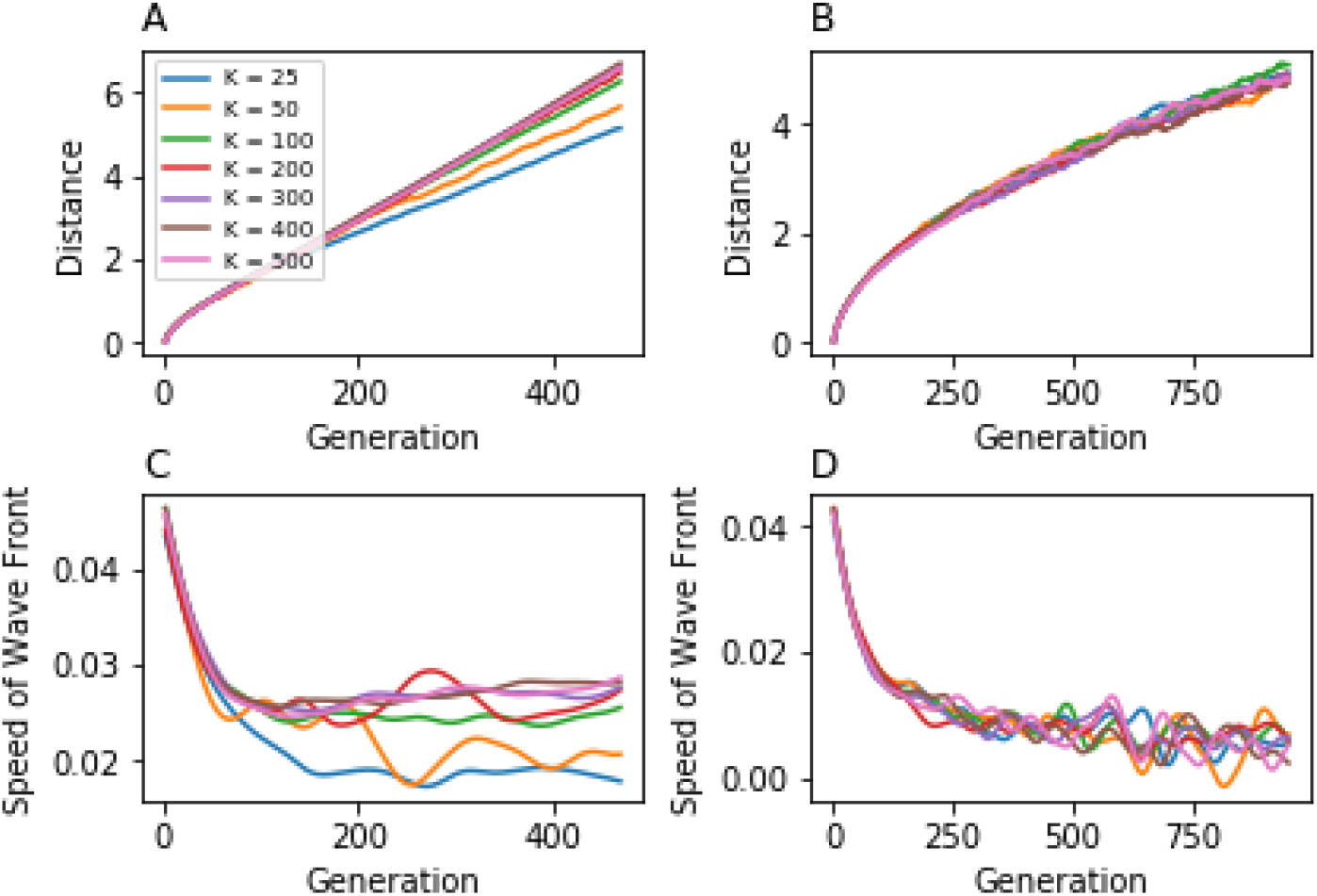
(A) Distances a selected allele (s = 0.05) was found from the point of origin *t* generations later. (B) Distances a neutral allele was found from point of origin. (C, D) Derivatives (speed) of the same corresponding simulations above. calculated with forward difference formula with *h* = 50 generations. *K* is representative of the population density.

**Figure 9:**
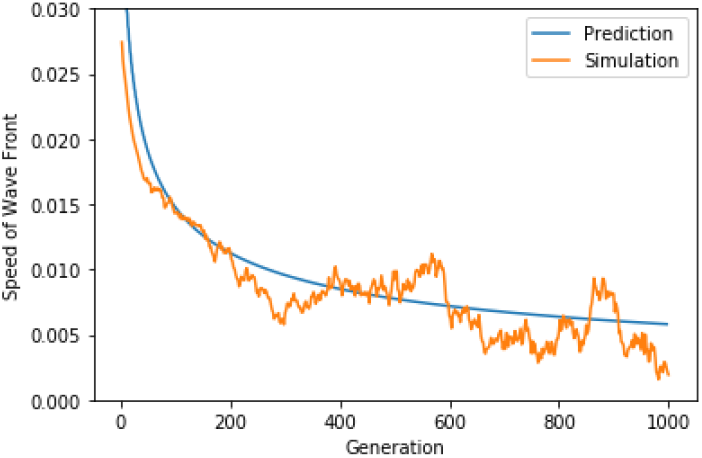
The average speed of simulated neutral allele and the prediction from our model.

To predict the speed of an advantageous allele we solved for equation 21 and let the results converge. We obtain a speed of about 0.023 for very small densities, and 0.03 for higher densities. Indeed, this is very close to what Figure 8 shows. Fisher’s equation evaluates to a speed of 0.0316 which is also close to what we see in our simulations. Zooming in and using boxplots to represent the range of speeds seen at each density helps portray some valuable insights (Figure 10). We can see that our model slightly overestimates the results from the simulations and seems to converge to Fisher’s prediction. This is good, as fisher did assume infinite population size (or rather density) so it is natural to view his results as the absolute maximum speed the allele could travel under infinite population densities. We can also see that our model predictions follow the shape seen in the simulations.

**Figure 10:**
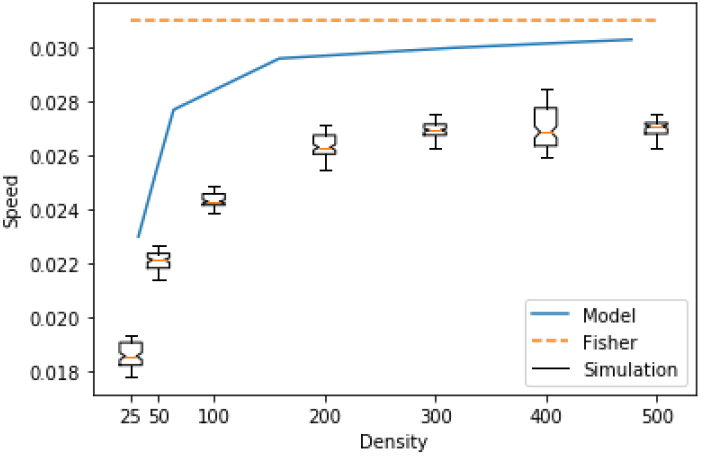
Box plots are the results from the simulation. Blue line represents the prediction from our model. The orange line represents Fisher’s prediction.

To truly see the difference between population size and density, we run several simulations in which population density remains constant while population size varied (Figure 11). This is accomplished by varying the geographical area a population occupies as observed from equation 1. Even though population size and population density can be very highly correlated, the speed at which new advantageous alleles travel depends only upon population density and not population size. Population size has no effect on the speed of the wave of advance. We argue that it also plays a minimal role in the relative strengths of selection and drift.

**Figure 11:**
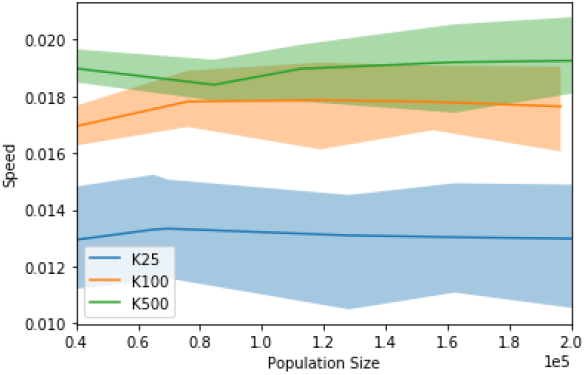
Wave speed versus population size for the population densities shown.

Figure 12 shows how the speed of advance is affected by selectional advantage. First note that increasing selective advantage clearly raises the curve, signifying an increase in the speed of the wave of advance. We also, again, see that the speed of the neutral allele is completely independent of population density, thus the flat line. The slightly advantageous alleles are also only slightly affected by population density while the selectively advantageous alleles of *s* = 0.05 and *s* = 0.1 clearly show the before noted pattern. That higher densities enable greater speed.

**Figure 12:**
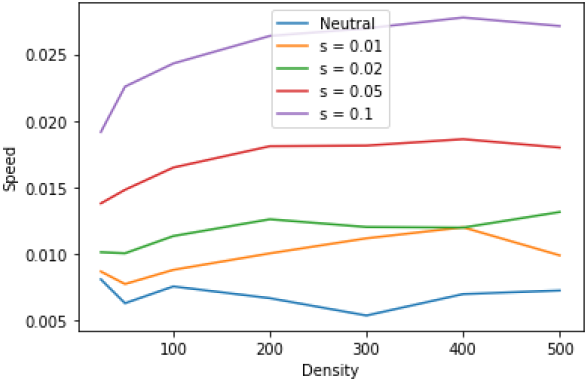
The velocity given population density, colored according to selectional advantage.

## 5. Discussion

We have shown that the rate at which new mutations saturate a population depends not on population size as might be expected, but rather on population density and geographical area. Now, it might be argued that population size is a function of population density and geographical area, so why not use population size as a compound variable to sit in for the other two parameters. We argue that if population density and geographical area are combined that there are several situations in which a model will fail to accurately portray a population.

Consider the expected time to fixation for an advantageous allele within a given population. If the population density were to increase, then both the population size would increase and the speed of the travelling wave of advance would increase. This will cause the new mutation to reach the outer edges of the population quicker and would reduce the expected time to fixation. However, say the population density remained constant while the geographical area increased. In this case the population size would still increase yet the speed of the wave of advance would not change. Since the geographical area is now larger it will take longer for the new mutation to reach the outer edges of the population and thus the time to fixation would increase. In both cases, the population size increases, but in one the time to fixation decreases while in the other it increases. There is no possible way this dynamic could be modelled with a single parameter. Thus, in this situation population size should be separated out into the two parameters of density and area.

We find it very interesting that the wave of advance of advantageous alleles travels at a constant rate. While it may seem intuitive, it could be extremely useful to know that any allele found expanding at a constant rate (i.e., radius of coverage growing linearly while area of coverage is growing quadratically), must be an advantageous allele. It also implies that species or alleles expanding at an ever-increasing rate are adapting must either be entering more suitable habitats thus allowing the population density to increase, or they are adapting to the environment in some way. These adaptations could either be increasing the selective advantage of their genome, or they could be increasing the dispersal distance as might be the case with the cane toads in Australia (Phillips et al., 2006, Urban et al., 2008).

The idea that the spread of neutral mutations is not affected by population density or population size, but only by the geographic area that the population covers is also very interesting. The only parameter needed to model these waves of advance was the dispersal distance. We note that population size must still play a role in the probability of fixation and in expectation for the time to fixation. But since an allele cannot fix until it has had the opportunity to reach the far edges of the geographic boundary the total geographic area of the species must also play a role.

We believe that as more and more species enter the endangered zone, that it will be crucial to understand how sparse populations differ from dense population more than we currently do. We recognize that the total population size determines the number of new mutations, but that it is the population density that determines the overall fate for each individual mutation. We feel it is important to continue this research, including an investigation into formulae for the expectation of fixation and time to fixation which incorporate population density and geographical area rather than population size. For future work it may also be interesting to explore the possibility of predicting the selective advantage of an allele based on the geographic spread. Smith et al. (2018) is one of many methods that estimate the age of an allele. A survey or sampling could be used to estimate the spread of an allele. Knowing the spread and time, it is easy to calculate the speed of the advance. If the population density is known then the selective advantage can be determined, and vice versa, if the selective coefficient is known then the population density can be estimated. Pockets of higher or lower density regions could be found when advantageous alleles travel through those regions at a different rate than expected.

## 6. Acknowledgement

The staff at the Research Computing Center at the Florida State University helped us to run our simulations smoothly on the high performance computing cluster.

